# *Vibrio natriegens*, a new genomic powerhouse

**DOI:** 10.1101/058487

**Authors:** Henry H. Lee, Nili Ostrov, Brandon G. Wong, Michaela A. Gold, Ahmad S. Khalil, George M. Church

## Abstract

Recombinant DNA technology has revolutionized biomedical research with continual innovations advancing the speed and throughput of molecular biology. Nearly all these tools, however, are reliant on *Escherichia coli* as a host organism, and its lengthy growth rate increasingly dominates experimental time. Here we report the development of *Vibrio natriegens*, a free-living bacteria with the fastest generation time known, into a genetically tractable host organism. We systematically characterize its growth properties to establish basic laboratory culturing conditions. We provide the first complete *Vibrio natriegens* genome, consisting of two chromosomes of 3,248,023 bp and 1,927,310 bp that together encode 4,578 open reading frames. We reveal genetic tools and techniques for working with *Vibrio natriegens*. These foundational resources will usher in an era of advanced genomics to accelerate biological, biotechnological, and medical discoveries.

---

Numerous recombinant DNA techniques have been developed to advance rapid and large-scale genomics (Litcofsky et al. 2012; H. H. Wang et al. 2009; Cohen et al. 1973; Gibson et al. 2009; Li and Elledge 2007; Engler et al. 2009). Nearly all of these techniques leverage the most rapidly growing model organism, *Escherichia coli*. Since most routine tasks, such as cloning, require the repeated growth of bacterial cultures, up to 90% of experimental time is allocated to *Escherichia coli* growth. A faster growing bacterial host would be highly desirable given the protracted and iterative nature of biological research.

*Vibrio natriegens* (previously *Pseudomonas natriegens* and *Beneckea natriegens*) is a Gram-negative, non-pathogenic marine bacterium isolated from salt marshes (Payne, Eagon, and Williams 1961). It is purported to be one of the fastest growing organisms known with a generation time between 7 to 10 minutes (Eagon 1962; Maida et al. 2013), Surprisingly, however, only few studies of *Vibrio natriegens* exist to date; little is known about the genetics that underlie its record setting replication rate and no tools or techniques have been described for working with this organism. We report the development of genetic methods for *Vibrio natriegens*, advancing the requisite tools to fully unravel the biology underlying its rapid growth and allowing its development as a superior alternative to *Escherichia coli*.

We started by investigating the conditions for stable culturing of *Vibrio natriegens*. We screened for readily-made, salt-rich growth media capable of supporting rapid and consistent growth. We found Lysogeny Broth supplemented with 1.5% Ocean Salts (LBO) or with 3% sodium chloride (LB3) to be the most robust growth media (Fig. 1a). We settled on LB3 as a standardized growth media due to the simplicity and accessibility of its formulation (Atkinson, M.J., and Bingham, C. 1997).

**Fig 1.**
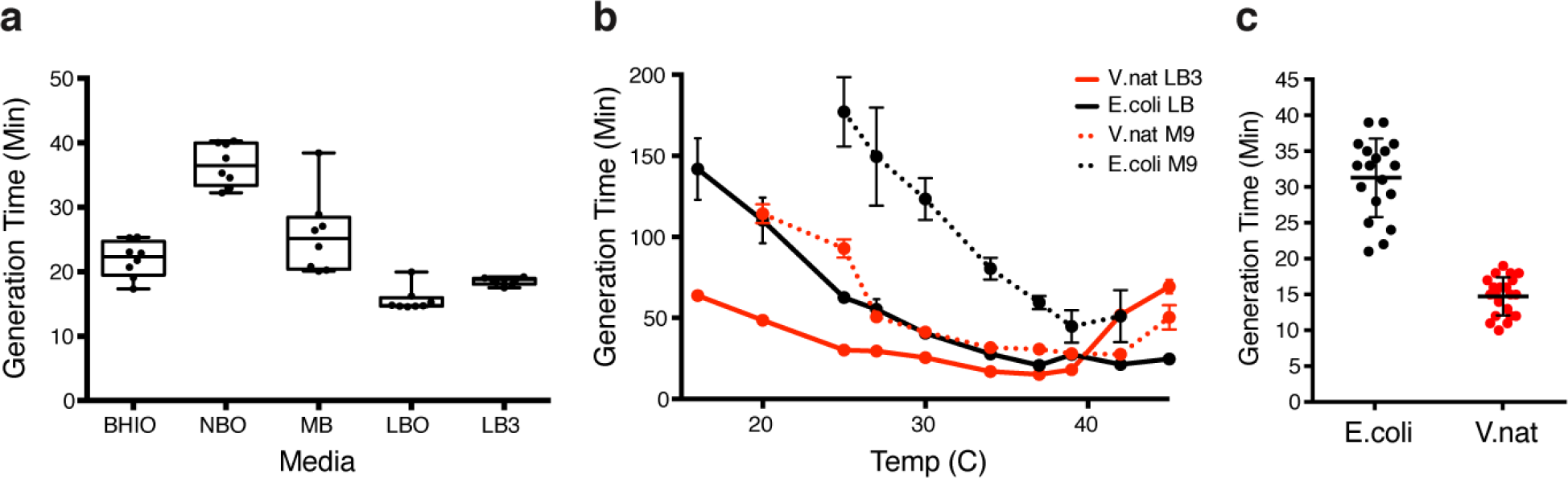
Characterization of *Vibrio natriegens* generation time in various conditions. (a) Bulk growth measurements of *Vibrio natriegens* growth in rich media supplemented with Ocean Salts: Brain Heart Infusion (BHIO), Nutrient Broth (NBO), Marine Broth (MB), Lysogeny Broth (LBO). LB3 is Lysogeny Broth with 3% (w/v) final of NaCl. Data shown are mean±SD (N=8). (b) Bulk growth measurements of *Escherichia coli* and *Vibrio natriegens* grown in LB and LB3, respectively, across a wide range of temperatures. Data shown are mean±SD (N=24) (c) Single-cell growth rate measurement at 37°C of *Escherichia coli* in LB and *Vibrio natriegens* in LB3. Data shown are mean±SD (N≥12).

We found that *Vibrio natriegens* outpaced *Escherichia coli* in both rich and minimal media at all tested temperatures under 42°C, with the fastest growth observed at 37°C (Fig. 1b). Specifically, *Vibrio natriegens* grew 1.4 to 2.2 times faster than *Escherichia coli* in rich media, and 1.6 to 3.9 times faster in minimal media supplemented with glucose. It is interesting to note that *Vibrio natriegens* grows robustly on sucrose, a economically and environmentally advantageous feedstock that the majority of current industrial *Escherichia coli* strains cannot utilize (Fig. S1) (Bruschi et al. 2012; Sabri, Nielsen, and Vickers 2013; Arifin et al. 2011).

To eliminate potential sources of error related to bulk measurements, we used time lapse single-cell microscopy to more accurately determine the relative generation times of *Vibrio natriegens* and *Escherichia coli* (Fig. 1c). Microfluidic chemostats were specifically designed to culture each of these organisms (Fig. S2, Supplementary Methods). At 37°C, *Vibrio natriegens*’ generation time was 14.8 minutes in LB3, 2.1 times faster than that of *Escherichia coli* in LB (31.3 minutes). These results confirm *Vibrio natriegens* as the fastest known free-living organism (Willardsen et al. 1978; Shimizu et al. 2002).

It is interesting to note that *Vibrio natriegens* is prone to lysis in salinities below that of ocean water despite having been isolated from estuarine regions, where salt water and fresh water mix (Fig. S6b). It is likely that this organism has adapted to an extreme ‘famine or feast’ lifestyle where ocean-like salinities exist for only short periods and highly rapid growth is thus under strong selection.

While cursory surveys can be performed on draft genomes, a fully annotated closed genome provides a solid foundation for further genetic investigation. We report *de novo* genome assembly of two closed circular chromosomes of 3.24 Mb (chr1) and 1.92 Mb (chr2), enabled by long-read SMRT sequencing (Supplementary Methods). The total genome size is approximately 5.17 Mb, over 0.5 Mb larger than the *Escherichia coli* genome (Fig. 2a). To our knowledge, this is the first report of a finished *Vibrio natriegens* genome allowing for an accurate accounting of all genes and their spatial organization, greatly improving upon previously reported draft genomes fragmented into multiple contigs (Z. Wang et al. 2013; Maida et al. 2013) (Table S1).

**Fig 2.**
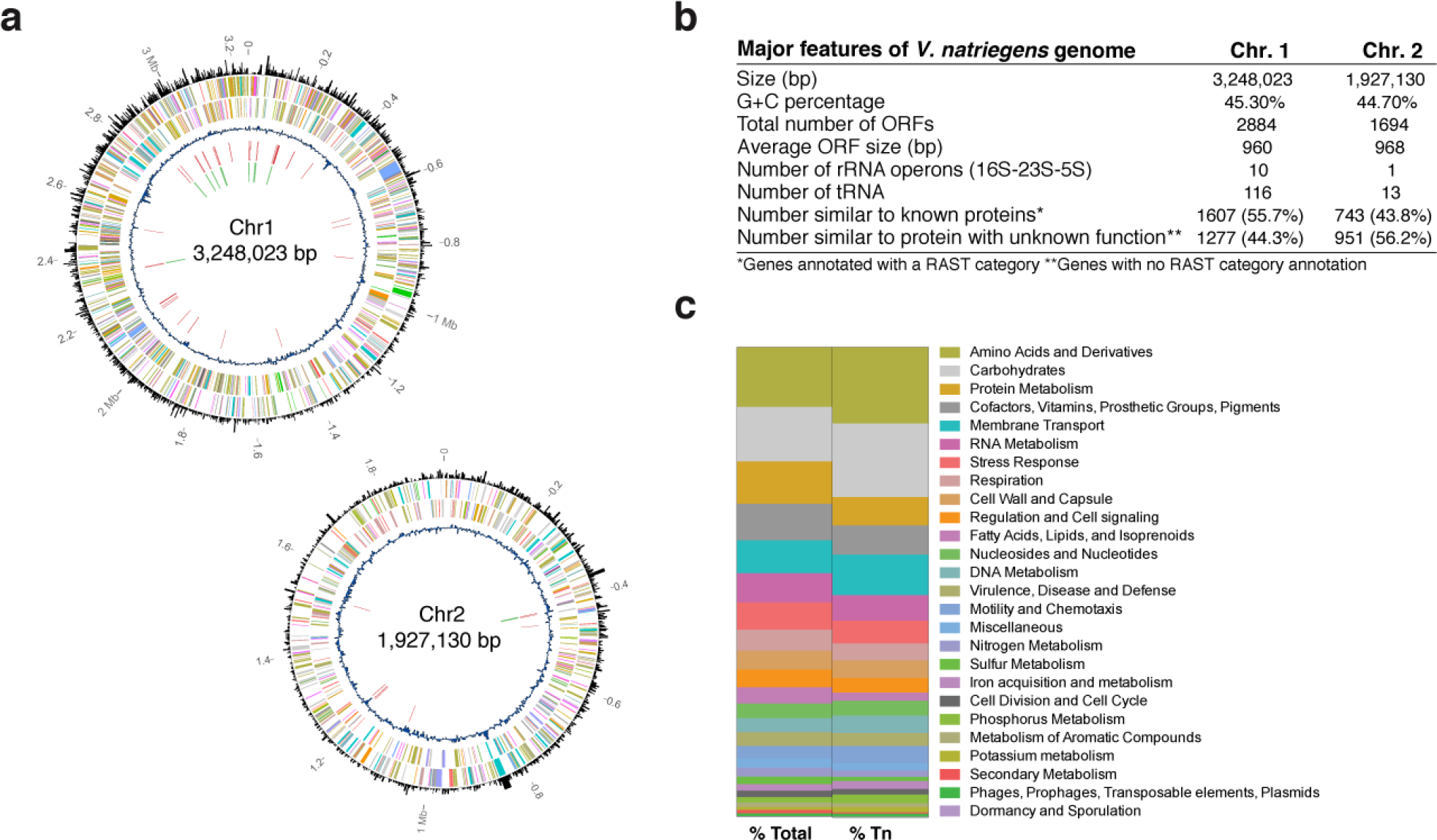
*Vibrio natriegens* genome. (a) The two circular chromosomes are depicted. From outside inward: Outermost circle indicates the number of transposons inserted in each gene. The next two circles represent protein-coding regions on the plus and minus strand, respectively, and are color coded by function as in panel c. Fourth circle represents G+C content relative to mean G+C content of the respective chromosome using a sliding window of 3,000 bp. Fifth circle shows tRNA genes. Sixth (innermost) circle shows rRNA genes. (b) Major features of *Vibrio natriegens* genome. (c) Percentage of genes annotated for each RAST category for the whole genome (Total) and for one or more insertions in the transposon mutant library (Tn).

We annotated the genome using the Rapid Annotation using Subsystem Technology (RAST) server (Fig. 2b) (Overbeek et al. 2013). Annotations revealed 11 rRNA operons, significantly more than *Escherichia coli* MG1655 and *Vibrio cholerae* El Tor N16961 with 7 and 8 rRNA operons, respectively (Heidelberg et al. 2000; Keseler et al. 2013). It is worth noting that previous work has implicated that a high number of rRNA operons contributes to the rapid growth rate of *Vibrio natriegens* (Aiyar, Gaal, and Gourse 2002). *Vibrio natriegens* carries 129 tRNA genes, 30 more than *Escherichia coli* MG1655 and 31 more than *Vibrio cholerae* El Tor N16961. We further analyzed the codon usage for this organism (Table S2). Given the growing interest in using orthogonal tRNA-aa pairs for expression of modified proteins, it is interesting to note only 706 instances would be required to re-assign the rarest codon genome-wide (UAG) (Lajoie et al. 2013). Notably, a preponderance of genes are associated with metabolic functions (Fig. 2c). It is intriguing to consider that formulation of specialized growth media along with highly oxygenated incubation conditions may uncover even faster growth rates for *Hbrio natriegens (Maida et al. 2013)*.

While DNA sequencing and genome analyses can yield valuable insights, experimentation is invaluable for deciphering biological functions *in vivo*. Modern genetic studies fundamentally depend on methods for introduction of recombinant DNA into the host, with electroporation or conjugation being the two most widely used. Developing DNA transformation methods for a largely unknown organism poses significant challenges. We describe our strategy for tackling these challenges and present initial results towards establishing tools for genetic studies in *Vibrio natriegens*.

No DNA transformation protocols or compatible plasmids have been described to date for *Vibrio natriegens*. Development of a robust transformation protocol presents a causality dilemma, since identifying a plasmid requires transformation, and development of a transformation protocol require a replicating plasmid. Without guiding information on the important properties or criteria to replicate a plasmid in *Vibrio natriegens*, we looked for plasmids that were found to replicate in a range of diverse bacteria. Such a plasmid, we reasoned, would improve our chances at observing transformants from initial and unoptimized electroporation screens. We thus selected a plasmid based on the RSF1010 operon, which carries its own host independent DNA initiator protein and primase, and has been found to replicate in a number of Gram-negative and some Gram-positive bacteria (Jain, Aayushi, and Preeti 2013). To differentiate transformants from possible escapees from antibiotic selection (Katashkina et al. 2009; Lutz and Bujard 1997), we engineered the RSF1010 replicon to constitutively express GFP so that we could visually check for successful transformants. We also determined the minimal concentration of antibiotic useful for plasmid selection (Table S3).

We then developed and optimized a rapid transformation protocol for *Vibrio natriegens* capable of achieving transformation efficiencies up to 2×10^5^ CFU / μg using our pRSF-tetO-GFP replicon (9.6 kb). We optimized the concentration of sorbitol as the osmoprotectant, voltage settings, input concentration of DNA, recovery media, and recovery time (Fig. S3). Our optimized transformation protocol can be performed in under 2 hours using as little as 10 ng of plasmid DNA. When incubated at 37°C, single transformant colonies can be visualized and picked within 5 hours of plating. Moreover, up to 2 μg of plasmid can be extracted from a single colony growing for 5 hours in liquid LB3 culture at 37°C, 2.5 times more than *Escherichia coli* similarly cultured in LB (Fig. S4). Notably, competent cells can be stored at −80°C and thawed for use with less than 4-fold loss of transformation efficiency in as little as 30 minutes (Supplementary Methods). Further optimization may improve transformation efficiencies for *Vibrio natriegens* to levels that rival *Escherichia coli*.

While our RSF1010 replicon plasmid allowed us to determine an electrotransformation protocol, its large size makes routine cloning unwieldy. Thus, we set out to find a smaller plasmid that would replicate in *Hbrio natriegens*. We first searched in NCBI Refseq for naturally occurring plasmids isolated from *Vibrios* (27 out of 2,647 total plasmids at the time of this study). However, the specific subset of sequences corresponding to the replicons from these plasmids is unknown and their large size (median plasmid size of 13 kb) precluded their synthesis due to cost (Boeke et al. 2016). Instead, we turned to mining bacteriophage replicons, inspired by the adaptation of the replicative form (RF) of M13 coliphage to construct some of the first replicating plasmids in *Escherichia coli* (Messing et al. 1977).

Interestingly, the vibriophage CTX, often studied in the context of pathogenic *Vibrio cholerae*, shares a similar life cycle to M13 and the genes responsible for its propagation have been determined (Waldor and Mekalanos 1996). Transformation of *Vibrio natriegens* with the CTX-Km RF yielded transformants which suggests that the CTX replicon is compatible in this host (Fig. S5a). We constructed a new plasmid, pRST, by fusing the specific replication genes from CTX-Km RF to a *Escherichia coli* plasmid based on the conditionally replicating R6k origin, thus adding a low-copy shuttle vector to the list of available genetic tools for *Vibrio natriegens*. We have used this plasmid in combination with the previously described pRSF plasmid as a dual plasmid system in *Vibrio natriegens* for complex regulation of proteins and high-throughput manipulation of diverse DNA libraries (Data not shown).

Importantly, we found that CTX vibriophage infects *Vibrio natriegens* at a rate that is at least 100-fold less active than infection of *Vibrio cholerae* O395 (Supplementary Methods). Given this extremely low infectivity, and the fact that CTX virions are not found in high-titers in the environment, *Vibrio natriegens* is an unlikely host for the propagation of the phage even if released into the environment (Davis 2003) (Fig. S5b). Even when the CTX RF is directly electroporated into *Vibrio natriegens*, we could not detect the production of infective CTX viral particles, suggesting that CTX viral particles are either not produced or not functionally assembled in *vibrio natriegens* (Fig. S5c). Furthermore, we were unable to find proteins homologous to known toxin genes or pathogenicity islands in the V. natriegens genome. These phenotypic and genotypic tests support the Biosafety Level 1 (BSL-1) designation for *Vibrio natriegens* as a generally safe biological agent.

Understanding the genetic pathways that contribute to the remarkable growth rate of *Vibrio natriegens* will likely require the development of high-throughput genome-wide methods to assess gene function. Whole-genome transposon mutagenesis has been used to generate large scale mutant libraries which when paired with high-throughput readout of their insertion position creates a powerful method for functional genomic screens (Gerdes et al. 2003; van Opijnen and Camilli 2010)

To perform whole-genome transposon mutagenesis in *Vibrio natriegens*, we first constructed a conjugative suicide vector based on the mariner transposase, previously used to mutagenize *Vibrio cholerae*. We then engineered the transposon mosaic end to permit high-throughput mapping of these insertions via Tn-seq (Goodman et al. 2009; van Opijnen and Camilli 2010; Cameron, Urbach, and Mekalanos 2008). In order for *Escherichia coli* to serve as a donor for conjugation into *Vibrio natriegens*, we determined the optimal salinity for both organisms. Finally, we optimized the conjugation time to maximize the number of unique transconjugants while minimizing clonal amplification (Fig. S6).

Our transposon mutagenesis protocol generated a library of 8.6×10^5^ *Vibrio natriegens* transconjugants. Significantly, over 97% of all Tn-Seq DNA reads fragments were successfully mapped to our reference genome, validating its accuracy. We found 4,562 unique insertions sites which were equally distributed between the two chromosomes (Fig. 2c). However, while 47% of all genes were mutagenized at least once, only 23.5% of all genes were found to carry two or more insertions, significantly lower than the coverage obtained when we applied this system to *Escherichia coli* (Table S4). Other transposon systems did not increase the number of insertions in the *vibrio natriegens* genome (Data not shown). A better understanding of the molecular mechanisms underlying transposition may be required to improve mutagenesis rates in *Vibrio natriegens*. This is the first genome-wide library available for high-throughput genetic screening in *Vibrio natriegens*.

Our transposon library can be used in diverse screens to isolate mutants with desirable phenotypes. Considering its already remarkable growth rate, we asked whether it would be possible to obtain *Vibrio natriegens* mutants capable of even faster growth. To do this, we mixed wild type and mutants in equal amounts and enriched for the fastest growing cells by culturing at a dilution rate that exceeded the maximum wild type growth rate. We determined the relative growth rates of mutant and wild-type cells by sampling the culture over time and discriminating the two populations via selective plating. Under these conditions, we were unable to identify mutant growing faster than wild type (Supplementary Methods). Relaxing this extreme growth selection may reveal mutants with subtle phenotypes worthy of further experimentation.

Unlike transposon mutagenesis, CRISPRi is capable of targeted gene inhibition but requires a genetic system capable of controlled expression with a measurable phenotype (Qi et al. 2013). To develop a CRISPRi system in *Vibrio natriegens*, we first established that the commonly used lactose and arabinose induction systems were operable, and characterized their dynamic ranges using GFP (Fig. S7) (Jacob and Monod 1961; Schleif 2000). We placed dCas9 under the control of arabinose promoter and the guide RNA under the control of the constitutive promoter J23100. Next, we used our transposon system to genomically integrate a constitutively expressed GFP construct. Using this engineered reporter strain, we show that inducing dCas9 in the presence of guide RNAs significantly inhibits chromosomal expression of GFP. Consistent with previous studies, we found stronger inhibition when using a guide RNA that targets the nontemplate strand (Fig. S8) (Larson et al. 2013). Further work is now ongoing to scale this promising tool for genome-wide perturbations (Peters et al. 2016).

Although we can now easily sequence an innumerable number of bacteria genomes, deeper understanding of cellular biology is still limited to a small number of highly studied species. Given the dominance of *Escherichia coli* as a genetic platform, it is not surprising that non-model bacteria remain largely genetically intractable. Development of host-agnostic genetic technologies and the comprehensive organization of these resources will be required to unlock novel and desirable phenotypes from diverse bacteria.

Taken together, this work introduces genetic tools towards developing *Vibrio natriegens* into a new genomic powerhouse. We provide the first complete genome sequence, standardize cultivation techniques and demonstrate its superior growth over *Escherichia coli* in a wide range of conditions. With genome mutagenesis, genetic screening, and targeted gene repression now possible, deeper investigations to uncover the genetic determinants of its unique growth properties will now be possible. These foundational resources will facilitate the advancement of *Vibrio natriegens* as faster and safe alternative to *Escherichia coli*.

## Acknowledgements

Henry Lee would like to acknowledge Javier Fernández Juárez, Reza Kalhor, Jun Teramoto, Nikolai Eroshenko, Michael Mee, Andrew Camilli, Jane Landolin, Tom Ellis, Anik Debnath, Alex Ng, Matthieu Landon, Harris Wang, and John Aach for helpful comments and discussions. Ahmad Khalil would additionally like to acknowledge Christopher Mancuso and Madeleine Joung. We thank Lyubov Golubeva for the pRSF plasmid, Brigid Davis and Matthew Waldor for *Vibrio cholerae* strains O395, BAH-2, and pCTX-Km and pCTX-Ap plasmids, Victor de Lorenzo for pBAM1 plasmid, D. Ewen Cameron and John Mekalanos for pTnFGL3 plasmid and Barry Wanner for BW29427 strain. This work was supported by Department of Energy Grant DE-FG02-02ER63445 and AWS Cloud Credits for Research program.

## Supplementary Figures

**Fig. S1.**
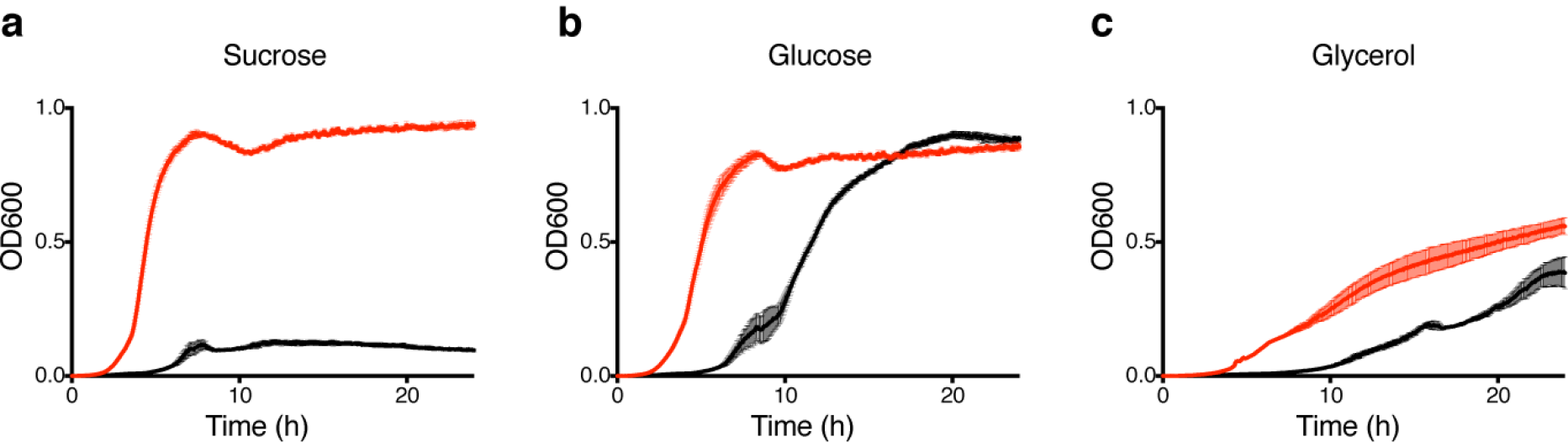
Bulk measurement of *Escherichia coli* (black) and *Vibrio natriegens* (red) growth in minimal media with the carbon sources as indicated.

**Fig. S2.**
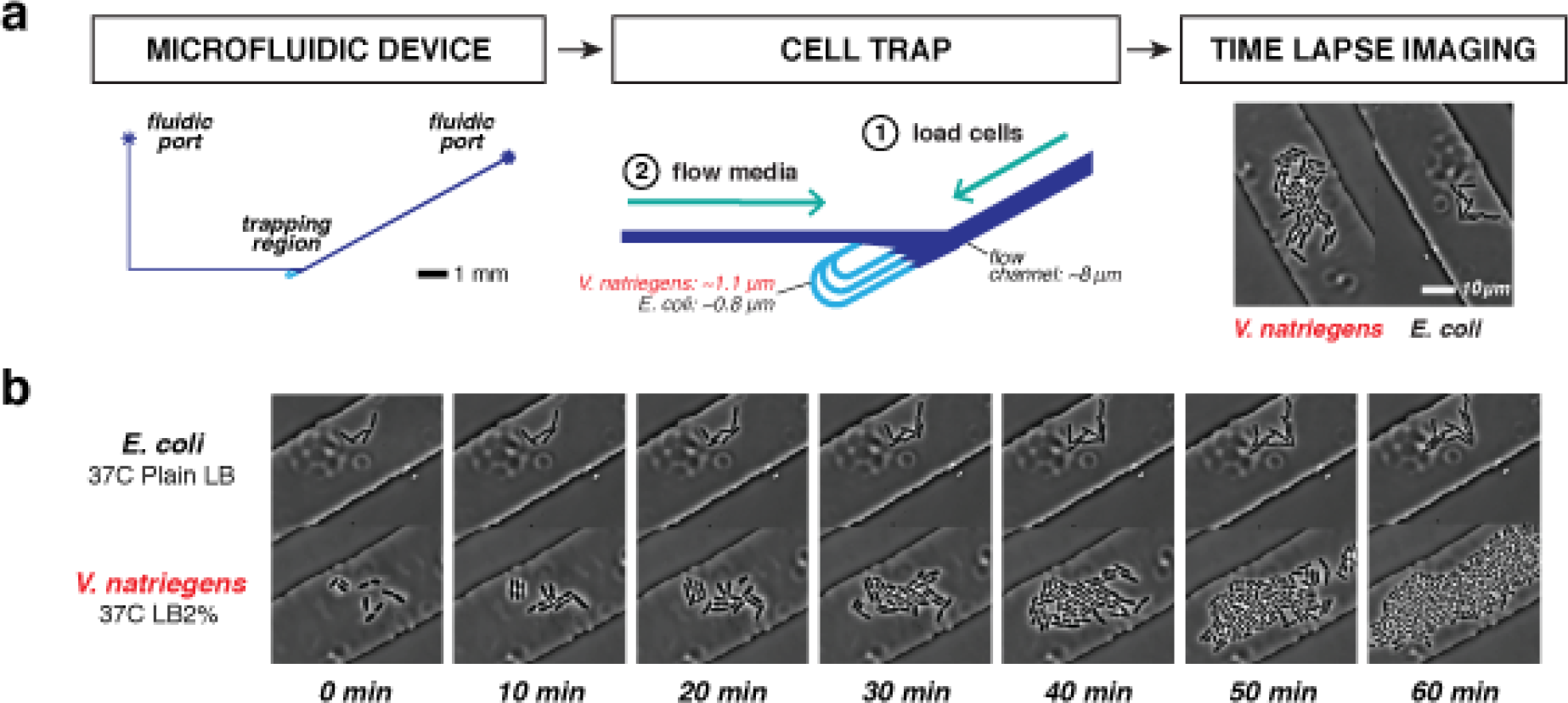
Single cell growth rate measurements. (a) Microfluidic chemostat devices keep cells growing in a monolayer. Two different devices, with altered cell trapping heights, were used to image *Vibrio natriegens* and *Escherichia coli*. (b) Side-by-side comparison of optimal growth conditions of each cell type.

**Fig. S3.**
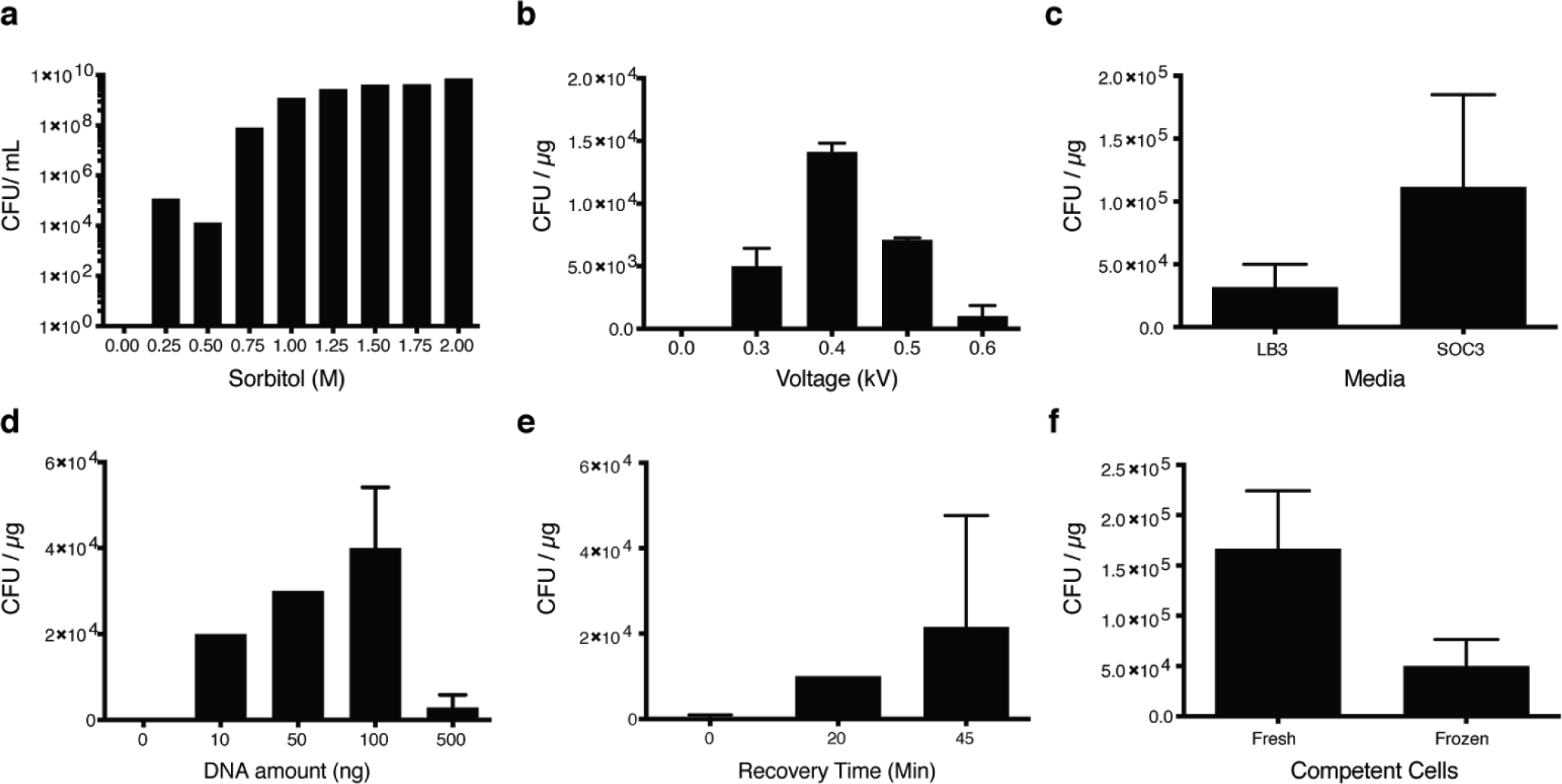
Optimization of parameters for *Vibrio natriegens* electroporation. (a) Cell viability in sorbitol, used as an osmoprotectant (representative data). Transformation efficiencies are shown for (b) Voltage. (c) Recovery media. (d) Amount of input plasmid DNA. (e) Recovery time. (d) Freshly prepared electrocompetent cells vs electrocompetent cells stored at −80°C. Unless otherwise indicated, transformations were performed using 50ng plasmid DNA with recovery time of 45min in SOC3. Data are shown as mean±SD (N≥2).

**Fig. S4.**
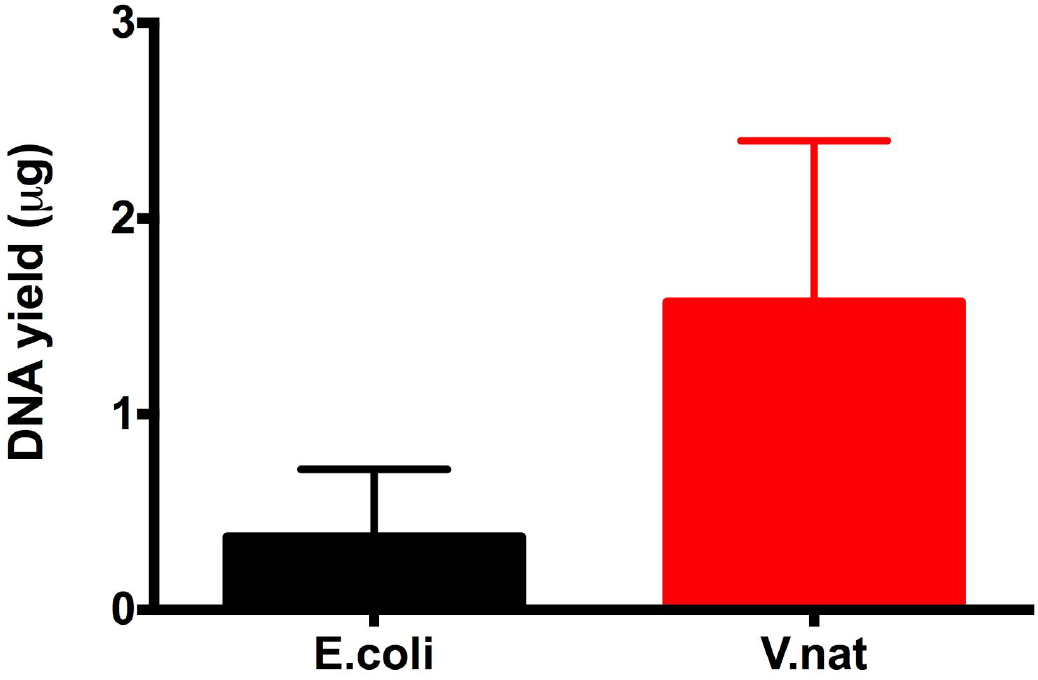
Rapid DNA production using *Vibrio natriegens*. Single colonies of *Vibrio natriegens* and *Escherichia coli* grown for 5 hours at 37°C in liquid LB3 or LB, respectively. Plasmid DNA is extracted from 3mL of each culture. Data are shown as mean±SD (N≥3).

**Fig. S5.**
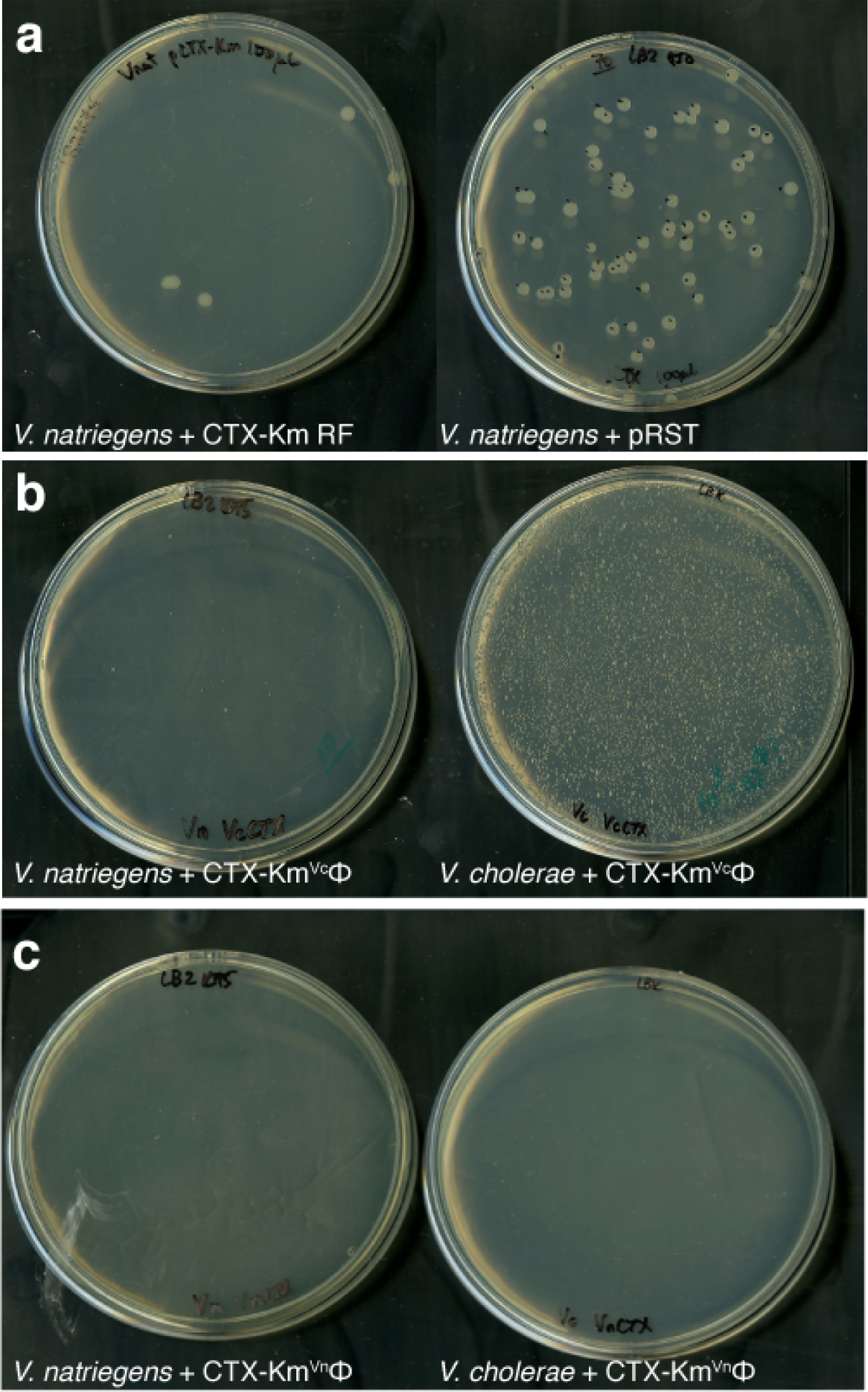
CTX replication and infectivity. (a) *Vibrio natriegens* transformants of CTX-Km RF (left) and pRST shuttle vector (right). (b) Transduction of *Vibrio natriegens* (left) and *Vibrio cholerae* O395 (right) by CTX-Km^Vc^Φ bacteriophage produced by *Vibrio cholerae* O395. (c) Transduction of *Vibrio natriegens* (left) and *Vibrio cholerae* O395 (right) by CTX-Km^Vn^Φ bacteriophage produced by *Vibrio natriegens*.

**Fig. S6.**
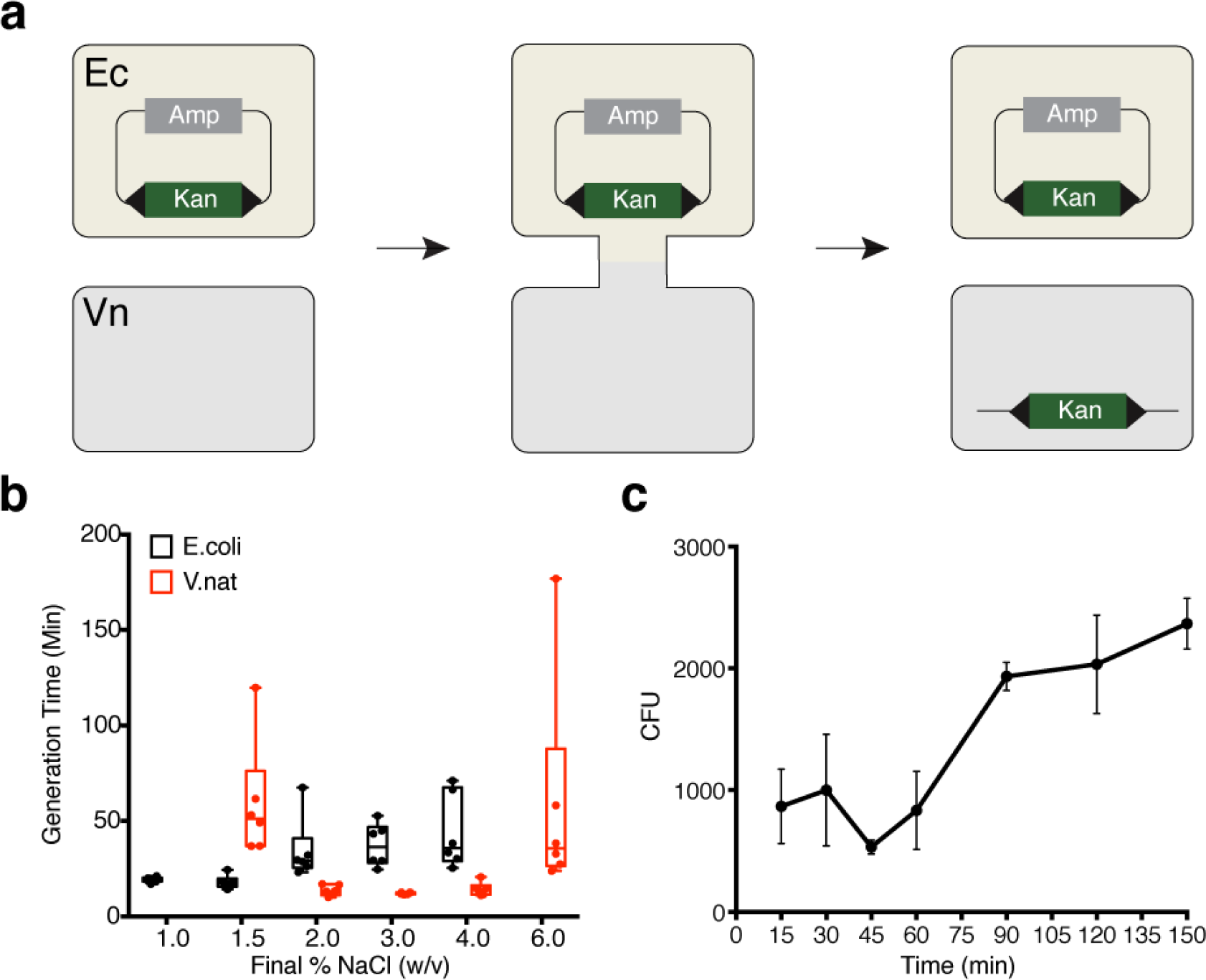
Determination of optimal conditions to conjugate a suicide vector for transposon mutagenesis. (a) Scheme for bi-parental conjugation between E. coli (Ec) and V. natriegens (Vn). (b) Cell viability as indicated by measurement of generation time in LB (corresponding to 1% NaCl) with increasing amount of salt. Note that V. Natriegens did not grow in LB (1% NaCl). (c) Number of transconjugants obtained as a function of conjugation time. Data are shown as mean±SD (N≥2).

**Fig. S7.**
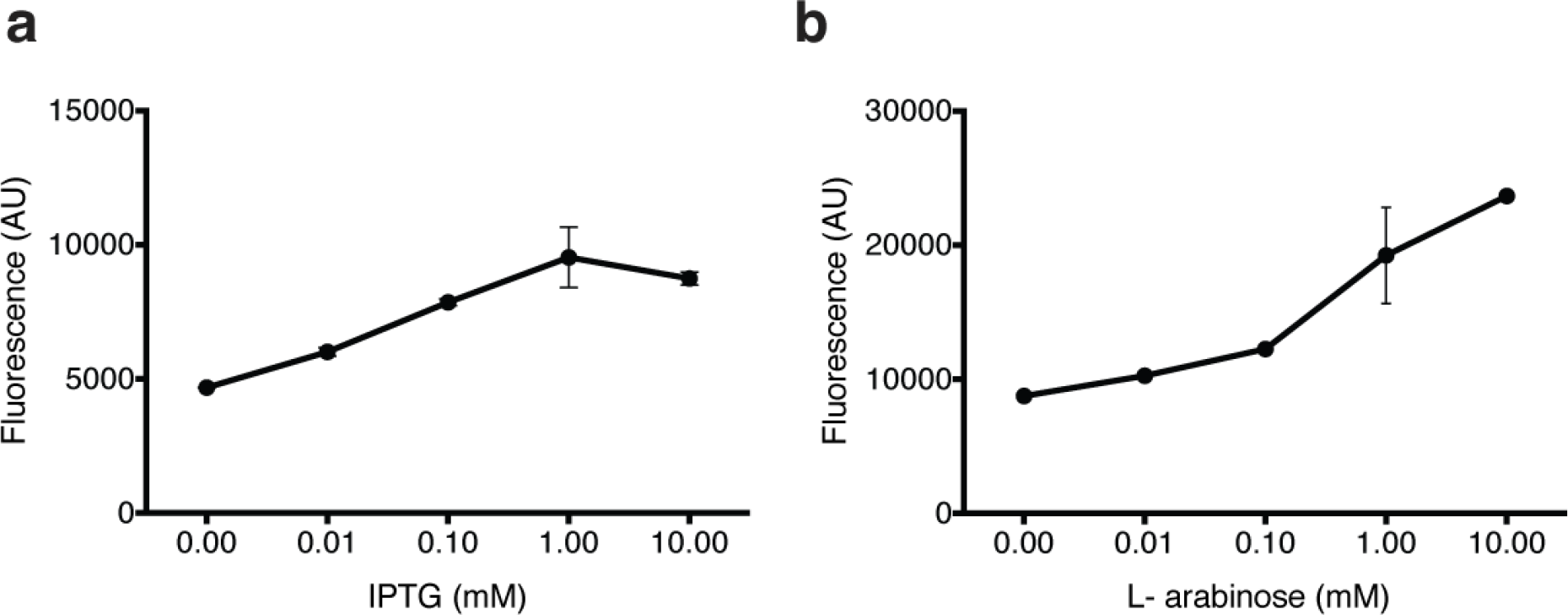
Titration of *Vibrio natriegens* induction systems. (a) Induction of the lactose promoter by IPTG. (b) Induction of the arabinose promoter by L-arabinose. Data are shown as mean±SD (N≥3).

**Fig. S8.**
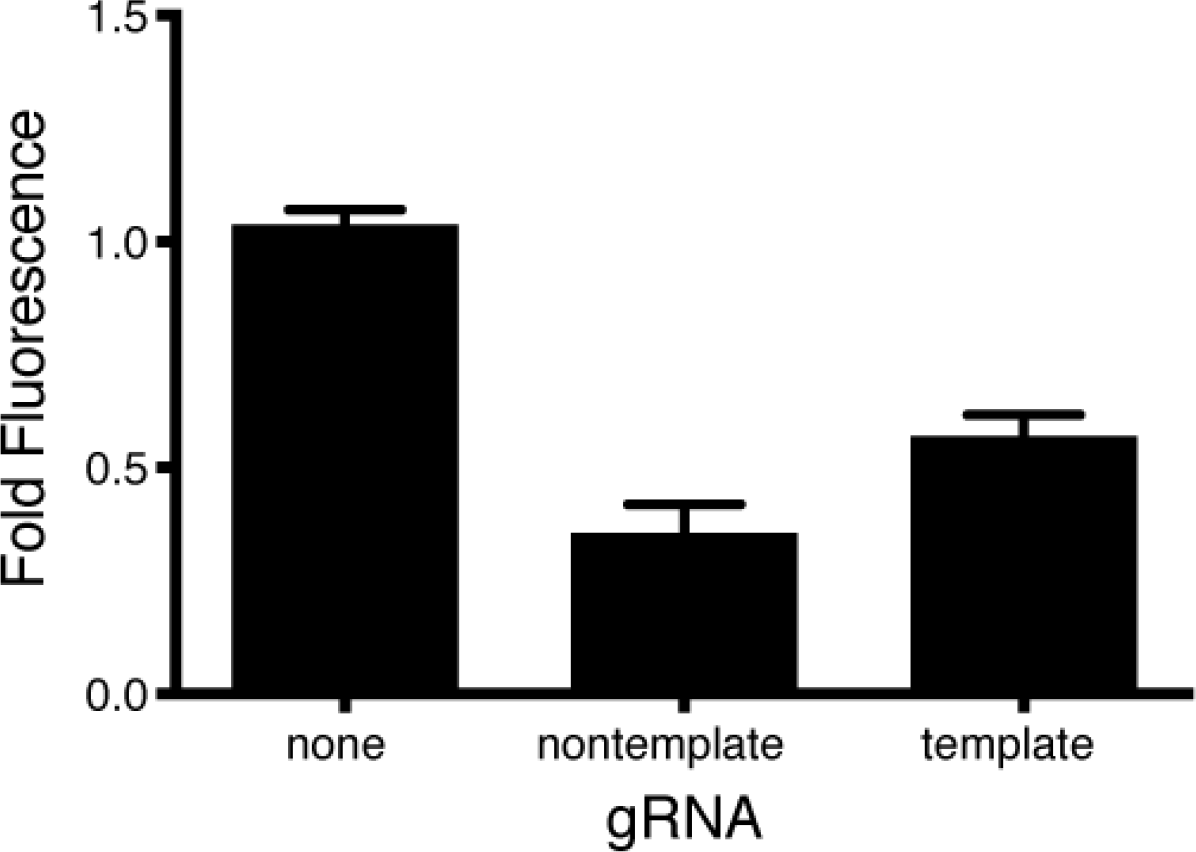
Targeted gene inhibition of chromosomally integrated GFP in *Vibrio natriegens* using dCas9. Guide RNA (gRNA) were designed to target the template or nontemplate strand of GFP. Data are shown as mean±SD (N≥3)

### Supplementary Tables

**Table S1.** Comparison of sequenced *Vibrio natriegens* genomes.

| Reference | Technology | Read Length<br>(bp) | Assembled<br>Genome Size (bp) | ORFs | Contigs | rRNA | tRNA |
| --- | --- | --- | --- | --- | --- | --- | --- |
| Maida et al. | Illumina | 101 | 5,200,362 | 4,788 | 173 | 14 | 71 |
| Wang et al. | Illumina | 2x250 | 5,131,685 | 4,587 | 45 | 12 | 124 |
| This work | PacBio | 4,407 (mean) | 5,175,333 | 4,578 | 2 (closed) | 11 | 129 |

**Table S2.** Codon usage of *Vibrio natriegens* genome.

| Codon | AA | Fraction | Frequency | Number | Codon | AA | Fraction | Frequency | Number |
| --- | --- | --- | --- | --- | --- | --- | --- | --- | --- |
| GCA | A | 0.297 | 25.499 | 37529 | CCA | P | 0.394 | 15.616 | 22983 |
| GCC | A | 0.16 | 13.753 | 20241 | CCC | P | 0.072 | 2.874 | 4230 |
| GCG | A | 0.279 | 23.949 | 35248 | CCG | P | 0.222 | 8.823 | 12985 |
| GCU | A | 0.264 | 22.657 | 33346 | CCU | P | 0.311 | 12.355 | 18184 |
| UGC | C | 0.336 | 3.508 | 5163 | CAA | Q | 0.59 | 25.924 | 38155 |
| UGU | C | 0.664 | 6.917 | 10181 | CAG | Q | 0.41 | 18.009 | 26506 |
| GAC | D | 0.395 | 21.56 | 31732 | AGA | R | 0.098 | 4.332 | 6376 |
| GAU | D | 0.605 | 33.05 | 48643 | AGG | R | 0.03 | 1.313 | 1933 |
| GAA | E | 0.651 | 42.243 | 62172 | CGA | R | 0.146 | 6.458 | 9505 |
| GAG | E | 0.349 | 22.632 | 33310 | CGC | R | 0.266 | 11.729 | 17262 |
| UUC | F | 0.408 | 16.819 | 24754 | CGG | R | 0.033 | 1.467 | 2159 |
| UUU | F | 0.592 | 24.364 | 35858 | CGU | R | 0.427 | 18.836 | 27723 |
| GGA | G | 0.122 | 8.499 | 12508 | AGC | S | 0.192 | 12.683 | 18666 |
| GGC | G | 0.324 | 22.604 | 33268 | AGU | S | 0.181 | 11.938 | 17570 |
| GGG | G | 0.098 | 6.814 | 10029 | UCA | S | 0.191 | 12.587 | 18526 |
| GGU | G | 0.457 | 31.901 | 46952 | UCC | S | 0.078 | 5.125 | 7543 |
| CAC | H | 0.498 | 11.015 | 16212 | UCG | S | 0.123 | 8.12 | 11951 |
| CAU | H | 0.502 | 11.121 | 16367 | UCU | S | 0.235 | 15.539 | 22870 |
| AUA | I | 0.102 | 6.423 | 9454 | ACA | T | 0.226 | 12.092 | 17797 |
| AUC | I | 0.415 | 25.998 | 38263 | ACC | T | 0.277 | 14.826 | 21820 |
| AUU | I | 0.483 | 30.288 | 44578 | ACG | T | 0.233 | 12.489 | 18381 |
| AAA | K | 0.685 | 35.536 | 52302 | ACU | T | 0.265 | 14.195 | 20892 |
| AAG | K | 0.315 | 16.332 | 24037 | GUA | V | 0.22 | 15.973 | 23509 |
| CUA | L | 0.13 | 13.323 | 19608 | GUC | V | 0.187 | 13.608 | 20028 |
| CUC | L | 0.089 | 9.132 | 13440 | GUG | V | 0.256 | 18.586 | 27354 |
| CUG | L | 0.223 | 22.871 | 33661 | GUU | V | 0.336 | 24.427 | 35952 |
| CUU | L | 0.181 | 18.499 | 27227 | UGG | W | 1 | 12.647 | 18614 |
| UUA | L | 0.195 | 19.94 | 29348 | UAC | Y | 0.561 | 16.918 | 24900 |
| UUG | L | 0.181 | 18.571 | 27333 | UAU | Y | 0.439 | 13.223 | 19461 |
| AUG | M | 1 | 26.992 | 39727 | UAA | * | 0.65 | 2.023 | 2977 |
| AAC | N | 0.561 | 23.211 | 34161 | UAG | * | 0.196 | 0.608 | 895 |
| AAU | N | 0.439 | 18.154 | 26719 | UGA | * | 0.154 | 0.48 | 706 |

**Table S3.** Concentrations of antibiotics used for plasmid selection in *Vibrio natriegens*.

| Antibiotic | µg/mL |
| --- | --- |
| Ampicillin | 100 |
| Kanamycin | 75 |
| Chloramphenicol | 1 |
| Spectinomycin | 100 |

**Table S4.** Number of genes, annotated by RAST category, for the whole genome and for the transposon library.

| RAST Category | All Annotated Genes |  | ≥1 Transposon Insertions |  |
| --- | --- | --- | --- | --- |
|  | # | % | # | % |
| Amino Acids and Derivatives | 366 | 12.72 | 209 | 16.21 |
| Carbohydrates | 334 | 11.61 | 202 | 15.67 |
| Protein Metabolism | 260 | 9.04 | 78 | 6.05 |
| Cofactors, Vitamins, Prosthetic Groups, Pigments | 224 | 7.79 | 80 | 6.21 |
| Membrane Transport | 202 | 7.02 | 111 | 8.61 |
| RNA Metabolism | 178 | 6.19 | 70 | 5.43 |
| Stress Response | 167 | 5.80 | 62 | 4.81 |
| Respiration | 129 | 4.48 | 46 | 3.57 |
| Cell Wall and Capsule | 116 | 4.03 | 49 | 3.80 |
| Regulation and Cell signaling | 109 | 3.79 | 40 | 3.10 |
| Fatty Acids, Lipids, and Isoprenoids | 99 | 3.44 | 22 | 1.71 |
| Nucleosides and Nucleotides | 89 | 3.09 | 41 | 3.18 |
| DNA Metabolism | 87 | 3.02 | 48 | 3.72 |
| Virulence, Disease and Defense | 84 | 2.92 | 35 | 2.72 |
| Motility and Chemotaxis | 78 | 2.71 | 48 | 3.72 |
| Miscellaneous | 55 | 1.91 | 19 | 1.47 |
| Nitrogen Metabolism | 55 | 1.91 | 18 | 1.40 |
| Sulfur Metabolism | 46 | 1.60 | 11 | 0.85 |
| Iron acquisition and metabolism | 40 | 1.39 | 22 | 1.71 |
| Cell Division and Cell Cycle | 39 | 1.36 | 15 | 1.16 |
| Phosphorus Metabolism | 34 | 1.18 | 24 | 1.86 |
| Metabolism of Aromatic Compounds | 26 | 0.90 | 10 | 0.78 |
| Potassium metabolism | 20 | 0.70 | 15 | 1.16 |
| Secondary Metabolism | 20 | 0.70 | 4 | 0.31 |
| Phages, Prophages, Transposable elements, Plasmids | 17 | 0.59 | 8 | 0.62 |
| Dormancy and Sporulation | 3 | 0.10 | 2 | 0.16 |
| Total | 2877 | 100.00 | 1289 | 100.00 |

## Materials and Methods

### Growth Media

Standardized growth media for *Vibrio natriegens* is named LB3 - Lysogeny Broth with 3% (w/v) final NaCl. We prepare this media by adding 20 grams of NaCl to 25 grams of LB Broth - Miller (Fisher BP9723-500). Rich media were formulated according to manufacturer instructions and supplemented with 1.5% final Ocean Salts (Aquarium System, Inc.) (w/v) to make high salt versions of Brain Heart Infusion (BHIO), Nutrient Broth (NBO), and Lysogeny Broth (LBO). No additional salts were added to Marine Broth (MB). Minimal M9 media was prepared according to manufacturer instruction. For culturing *Vibrio natriegens*, 2% (w/v) final sodium chloride was added to M9. Carbon sources were added as indicated to 0.4% (v/v). Unless otherwise indicated, *Vibrio natriegens* experiments were performed in LB3 media and *Escherichia coli* experiments were performed in LB media. SOC3 media is composed of 5 grams of yeast extract, 20 grams tryptone, 30 grams sodium chloride, 2.4 grams magnesium sulfate, and 0.4% (v/v) final glucose.

### Overnight culturing

An inoculation of −80°C frozen stock of *Vibrio natriegens* can reach stationary phase after 5 hours when incubated at 37°C. Prolonged overnight culturing (>15 hours) at 37°C can lead to an extended lag phase upon subculturing. Routine overnight culturing of *Vibrio natriegens* is performed for 8-15 hours at 37°C or 12-24 hours at room temperature. Unless otherwise indicated, *Escherichia coli* cells used in this study were K-12 subtype MG1655 unless otherwise indicated and cultured overnight (>10 hours) at 37°C. *Vibrio cholerae* O395 was cultured overnight (>10 hours) in LB at 30°C or 37°C in a rotator drum at 150rpm.

### Glycerol Stock

To prepare *Vibrio natriegens* cells for −80°C storage, an overnight culture of cells must be washed in fresh media before storing in glycerol. A culture was centrifuged for 1 minute at 20,000 rcf and the supernatant was removed. The cell pellet was resuspended in fresh LB3 media and glycerol was added to 20% final concentration. The stock is quickly vortexed and stored at −80°C. Note: unlike glycerol stocks of *Escherichia coli* for −80°C storage, neglecting the washing step prior to storing *Vibrio natriegens* cultures at −80°C can lead to an inability to revive the culture.

### Bulk measurements of generation time

Growth was measured by kinetic growth monitoring (Biotek H1, H4, or Eon plate reader) in 96-well plates with continuous orbital shaking and optical density measurement at 600nm taken every 2 minutes. Overnight cells were washed once in fresh growth media, then subcultured by at least 1:100 dilution. To assay *Vibrio natriegens* growth in different rich media, cells were cultured overnight from frozen stock in the specific rich media to be tested. To assay *Vibrio natriegens* and *Escherichia coli* growth in minimal media, cells were cultured overnight in LB3 and LB respectively, and subcultured in the appropriate test media. Generation times are calculated by linear regression of the log-transformed OD across at least 3 data points when growth was in exponential phase. To avoid specious determination of growth rates due to measurement noise, the minimal OD considered for analysis was maximized and the ODs were smoothed with a moving average window of 3 data points for conditions that are challenging for growth.

### Microfluidics device construction

Microfluidic devices were used as tools to measure and compare growth rates of *E. coli* and *V. natriegens* in several different growth conditions. In these devices, cells are grown in monolayer and segmented/tracked in high temporal resolution using time-lapse microscopy. The cells are constricted for imaging using previously described Tesla microchemostat device designs (Cookson et al. 2005; Stricker et al. 2008; Vega et al. 2012), in which cell traps have heights that match the diameters of the cells, minimizing movement and restricting growth in a monolayer. Different trapping heights of 0.8 μm and 1.1 μm were used for *E. coli* and *V. natriegens*, respectively.

The microfluidic devices were fabricated with polydimethylsiloxane (PDMS/Sylgard 184, Dow Corning) using standard soft lithographic methods (Duffy et al. 1998). Briefly, the microfluidic devices were fabricated by reverse molding from a silicon wafer patterned with two layers of photoresist (one for the cell trap, another for flow channels). First, the cell trap layer was fabricated by spin coating SU-8 2 (MicroChem Corp.) negative resist at 7000 RPM and 6800 RPM, for *E. coli* and *V. natriegens* respectively, and patterned using a high resolution photomask (CAD/Art Services, Inc.). Next, AZ4620 positive photoresist (Capitol Scientific, Inc.) was spun onto the silicon wafer and aligned with another photomask for fabrication of ~8 μm tall flow channels (same for both organisms). Reverse-molded PDMS devices were punched and bonded to No. 1.5 glass coverslips (Fisher Scientific), similar to previously described protocols (Duffy et al. 1998).

### Time-lapse microscopy and image analysis

Cells were diluted down to 0.1 OD_600_ from an overnight culture at optimal growth conditions and allowed to grow for an hour in the corresponding media conditions (e.g. temperature, salt concentration) before loading onto the device. Next, cells were loaded and grown on the device in the corresponding environmental conditions until the cell trap chambers filled. Temperature was maintained with a Controlled Environment Microscope Incubator (Nikon Instruments, Inc.). Media flow on device was maintained by a constant pressure of 5 psi over the course of the experiment after cell loading.

During the experiment, phase contrast images were acquired every minute with a 100x objective (Plan Apo Lambda 100X, NA 1.45) using an Eclipse Ti-E inverted microscope (Nikon Instruments, Inc.). Images were acquired using “Perfect Focus”, a motorized stage, and a Clara-E charge-coupled device (CCD) camera (Andor Technology). After the experiment, images were segmented using custom MATLAB (Mathworks, Natick, MA) software. Code available upon request.

### Preparation of electrocompetent *Vibrio natriegens*

*Vibrio natriegens* was grown overnight as indicated above. A subculture was prepared by inoculating an overnight culture, washed once in fresh LB3, at 1:100 dilution into fresh media. For example, 500μL of the overnight culture was pelleted by centrifugation for 1 minute at 20,000rcf, resuspended in 500μL fresh LB3 media and inoculated in 50mL of LB3 media. The culture was incubated at 37°C at 225rpm for 1 hour reaching OD ~0.4. Cells were then pelleted by centrifugation at 3500rpm for 5min at 4°C, and washed by resuspension in 1ml of cold 1M sorbitol followed by centrifugation at 20,000rcf for 1 minute at 4°C. The wash was repeated for a total of three times. The final cell pellet was resuspended in 250μL of 1M sorbitol. 50μL of concentrated cells were used per transformation. For long term storage, the concentrated cells were aliquoted in 50μL shots in chilled tubes, snap frozen in dry ice and ethanol, and stored in −80°C for future use. To transform, 50ng of plasmid DNA was added to the cells in 0.1mm cuvettes and electroporated using Biorad Gene Pulser electroporator at 0.4kV, 1kΩ, 25μF. Cells were recovered in 1mL LB3 or SOC3 media for 45 minutes at 37°C at 225rpm, and plated on selective media. Plates were incubated at least 6 hours at 37°C or at least 12 hours at room temperature.

### Plasmid constructions

Routine cloning was performed by PCR of desired DNA fragments, assembly with NEB Gibson Assembly or NEBuilder HiFi DNA Assembly, and propagation in *Escherichia coli* (Gibson and Daniel 2009) unless otherwise indicated. We used pRSF for the majority of our work since it carries all of its own replication machinery and should be minimally dependent on host factors (Katashkina et al. 2009). For our transformation optimizations, we constructed pRSF-pLtetO-gfp, which constitutively expresses GFP due to the absence of the tetR repressor in both *Escherichia coli* and *Vibrio natriegens*. We engineered our pRST shuttle plasmid by fusing the pCTX-Km replicon with the pir-dependent conditional replicon, R6k. To construct the conjugative suicide mariner transposon, we replaced the Tn5 transposase and Tn5 mosaic ends in pBAM1 with the mariner C9 transposase and the mariner mosaic ends from pTnFGL3 (Cameron, Urbach, and Mekalanos 2008; Martínez-García et al. 2011). Our payload, the transposon DNA, consisted solely of the minimal kanamycin resistance gene required for transconjugant selection. We next performed site-directed mutagenesis on both transposon mosaic ends to introduce an Mmel cut-site, producing the plasmid pMarC9 which is also based on the pir-dependent conditional replicon, R6k. We also constructed a transposon plasmid capable of integrating a constitutively expressing GFP cassette in the genome by inserting pLtetO-GFP with either kanamycin or spectinomycin in the transposon DNA. All plasmids carrying the R6k origin was found only to replicate in either BW29427 or EC100D pir^+^/pir-116 *Escherichia coli* cells. Induction systems were cloned onto the pRSF backbone. For the CRISPRi system, we utilized a single plasmid carrying both dCas9, the nuclease-null *Streptococcus pyogenes* cas9, and the guide RNA. The dCas9 was under the control of arabinose induction and the guide RNA was under control of the constitutive J23100 promoter.

### CTX vibriophage infection

*Vibrio cholerae* O395 carrying the replicative form of CTX, CTX-Km (kanamycin resistant) was cultured overnight in LB without selection in a rotator drum at 150rpm at 30°C. Virions were purified from cell-free supernatant (0.22μm filtered) of overnight cultures. Replicative forms were extracted from the cells by standard miniprep (Qiagen). To test infectivity of the virions, naive *Vibrio cholerae* O395 and *Vibrio natriegens* were subcultured 1:1000 in LB and LB3 respectively and mixed gently with approximately 10^6^ virions. After static incubation for 30 minutes at 30°C, the mixture was plated on selective media and incubated overnight for colony formation. Replicative forms were electroporated into host strains using described protocols.

### Genome sequencing, assembly, and annotation

*Vibrio natriegens* (ATCC 14048) was cultured for 24 hours in Nutrient Broth with 1.5% NaCl according to ATCC instructions. Genomic DNA was purified (Qiagen Puregene Yeast/Bact. Kit B) and sequenced on a Single Molecule Real Time (SMRT) Pacific Biosciences RS II system (University of Massachusetts Medical School Deep Sequencing Core) using 120 minute movies on 3 SMRTCells. SMRTanalysis v2.1 on Amazon Web Services was used to process and assemble the sequencing data. The mean read length, after default quality filtering, was 4,407bp. HGAP3 with default parameters was used to assemble the reads which yielded 2 contigs. The contigs were visualized with Gepard and manually closed (Krumsiek, Arnold, and Rattei 2007). The two closed chromosomes annotated using RAST under ID 691.14 (Aziz et al. 2008). The annotated genome is deposited in NCBI under Biosample SAMN03178087, GenBank CP009977 and CP009978. Codon usage was calculated using EMBOSS cusp.

### Construction of transposon mutant libraries

Our conjugative suicide mariner transposon plasmid was propagated in BW29427, an *Escherichia coli* with diaminopimelic acid (DAP) auxotrophy. BW29427 growth requires 300μM of DAP even when cultured in LB. Importantly, BW29427 does not grow when DAP is not supplemented, which simplifies counterselection of this host strain following biparental mating with *Vibrio natriegens*. To conjugate from *Escherichia coli* to *Vibrio natriegens*, strains were grown to OD 0.4 and equal volumes of each culture was spun down and resuspended in a minimal volume of LB2 (Lysogeny Broth with 2% (w/v) final of sodium chloride). The cell mixture is dispensed onto an LB2 plate with or without a 0.22 μm filter (Millipore), and incubated at 37°C for 60 minutes. The cells are harvested from the surface of the plate either by directly scraping or vortexing the filter in 1mL of LB3 media. The resulting cell resuspension is washed once in fresh LB3, resuspended to a final volume of 1mL, and plated on 245mm × 245mm kanamycin selective plates (Corning). Plates were incubated at 30°C for 12 hours to allow the formation of *Vibrio natriegens* colonies. BW29427 colonies were not detected on the resulting plate, suggesting that host counterselection was successful. Colonies were scraped from each plate with 3mL of LB3, gently vortexed, and stored as glycerol stock as previously described. A similar protocol was used to generate an *Escherichia coli* transposon mutant library, except LB was used as the media at all steps.

### Analysis of the transposon mutant library

Tn-seq was performed as previously described (van Opijnen and Camilli 2010). Briefly, genomic DNA was extracted (Qiagen DNeasy Blood & Tissue Kit), and digested with MmeI. To enrich for the fragment corresponding to the kanamycin transposon fragment, the digested genomic DNA was electrophoresed on a 1% TAE gel and an area of the gel corresponding to approximately 1.2kb was extracted. The resulting DNA fragment was sticky-end ligated to an adapter. PCR was used to selectively amplify the region around the transposon mosaic end and to add the required Illumina adapters. These amplicons were sequenced 1×50bp on a MiSeq. Since properly prepared amplicons contain 16 or 17bp of genomic DNA and 32 or 33bp of the ligated adapter, only those sequencing reads with the presence of the adapter were further analyzed. All adapters were trimming and the resulting genomic DNA sequences were aligned to the reference genome with Bowtie (Langmead et al. 2009).

### Growth screen for *Vibrio natriegens* mutants with hyper-growth phenotypes

Wild-type and mutant *Vibrio natriegens* cells were grown independently at 37°C until OD ~0.8. The cultures were mixed in equal volume and grown in a turbidostat at a dilution rate of 15mL/hr (d=6h^−1^) (Takahashi et al. 2015). For the selection, the dilution rate was increased to 17mL/hr. Samples of the culture was drawn at different ODs as it decayed. These samples were plated on permissive (LB3) and selective plates (LB3+kanamycin) to determine the ratio of wild-type cells to mutant cells, respectively. At OD ~0.1, only 1% of the population were kanamycin resistant. At OD ~0.04, no kanamycin resistant cells were detected.

### Arabinose and IPTG Induction assay

*Vibrio natriegens* carrying plasmid pRSF-pBAD-GFP or pRSF-pLacI-GFP were used for all induction assays. Overnight cultures were washed with LB3 media and diluted 1:1000 into selective LB3 media with varying concentration of IPTG or L-arabinose. OD600 and fluorescence were kinetically monitored in a microplate with orbital shaking at 37°C. Fluorescence after 7 hours of culturing is shown.

### Repression of chromosomally-encoded GFP with CRISPRi

We used our previously described transposon system to chromosomally integrated a cassette that constitutively expresses GFP. We transformed this engineered *Vibrio natriegens* strain with our CRISPRi plasmid carrying both dcas9 and GFP-targeting gRNA. To test the repression of the chromosomally-encoded GFP with CRISPRi, we subcultured the overnight cultures 1:1000 in fresh media supplemented with or without 1mM arabinose. We kinetically measured OD600 and fluorescence of each culture over 12 hours in a microplate with orbital shaking at 37°C. In these conditions, all cultures grew equivalently. Fold repression was calculated as the ratio of final fluorescence for each construct with or without the addition of arabinose.

### DNA yield

We transformed pRSF-pLtetO-gfp via electroporation into *Vscherichia coli* K-12 MG1655 and *Vibrio natriegens*. The *Escherichia coli* plate was incubated at 37°C and the *Vibrio natriegens* plate was incubated at room temperature for an equivalent time to yield approximately similar colony sizes. 3 colonies from each plate was picked and grown for 5 hours at 37°C in 3mL of selective liquid culture (LB for *Escherichia coli* and LB3 for *Vibrio natriegens*) at 225rpm. Plasmid DNA was extracted from 3mL of culture (Qiagen Plasmid Miniprep Kit).

## References

Aiyar, Sarah E., Tamas Gaal, and Richard L. Gourse. 2002. “rRNA Promoter Activity in the Fast-Growing Bacterium Vibrio Natriegens.” Journal of Bacteriology 184 (5): 1349–58.

Arifin, Yalun, Arifin Yalun, Sabri Suriana, Sugiarto Haryadi, Jens O. Krömer, Claudia E. Vickers, and Lars K. Nielsen. 2011. “Deletion of cscR in Escherichia Coli W Improves Growth and Poly-3-Hydroxybutyrate (PHB) Production from Sucrose in Fed Batch Culture.” Journal of Biotechnology 156 (4): 275–78.

Atkinson, M.J., and Bingham, C. 1997. “Elemental Composition of Commercial Seasalts.” Journal of Aquariculture and Aquatic Sciences VIII (2): 39–43.

Boeke, Jef D., George Church, Andrew Hessel, Nancy J. Kelley, Adam Arkin, Yizhi Cai, Rob Carlson, et al. 2016. “The Genome Project-Write.” Science, June. doi:10.1126/science.aaf6850.

Bruschi, Michele, Simon J. Boyes, Haryadi Sugiarto, Lars K. Nielsen, and Claudia E. Vickers. 2012. “A Transferable Sucrose Utilization Approach for Non-Sucrose-Utilizing Escherichia Coli Strains.” Biotechnology Advances 30 (5): 1001–10.

Cameron, D. Ewen, Jonathan M. Urbach, and John J. Mekalanos. 2008. “A Defined Transposon Mutant Library and Its Use in Identifying Motility Genes in Vibrio Cholerae.” Proceedings of the National Academy of Sciences of the United States of America 105 (25): 8736–41.

Cohen, S. N., A. C. Chang, H. W. Boyer, and R. B. Helling. 1973. “Construction of Biologically Functional Bacterial Plasmids in Vitro.” Proceedings of the National Academy of Sciences of the United States of America 70 (11): 3240–44.

Davis, B. 2003. “Filamentous Phages Linked to Virulence of Vibrio Cholerae.” Current Opinion in Microbiology 6 (1): 35–42.

Eagon, R. G. 1962. “Pseudomonas Natriegens, a Marine Bacterium with a Generation Time of Less than 10 Minutes.” Journal of Bacteriology 83 (April): 736–37.

Engler, Carola, Ramona Gruetzner, Romy Kandzia, and Sylvestre Marillonnet. 2009. “Golden Gate Shuffling: A One-Pot DNA Shuffling Method Based on Type IIs Restriction Enzymes.” PloS One 4 (5): e5553.

Gerdes, S. Y., M. D. Scholle, J. W. Campbell, G. Balázsi, E. Ravasz, M. D. Daugherty, A. L. Somera, et al. 2003. “Experimental Determination and System Level Analysis of Essential Genes in Escherichia Coli MG1655.” Journal of Bacteriology 185 (19): 5673–84.

Gibson, Daniel G., Young Lei, Chuang Ray-Yuan, J. Craig Venter, Clyde A. Hutchison, and Hamilton O. Smith. 2009. “Enzymatic Assembly of DNA Molecules up to Several Hundred Kilobases.” Nature Methods 6 (5): 343–45.

Goodman, Andrew L., Nathan P. McNulty, Yue Zhao, Douglas Leip, Robi D. Mitra, Catherine A. Lozupone, Rob Knight, and Jeffrey I. Gordon. 2009. “Identifying Genetic Determinants Needed to Establish a Human Gut Symbiont in Its Habitat.” Cell Host & Microbe 6 (3): 279–89.

Heidelberg, J. F., J. A. Eisen, W. C. Nelson, R. A. Clayton, M. L. Gwinn, R. J. Dodson, D. H. Haft, et al. 2000. “DNA Sequence of Both Chromosomes of the Cholera Pathogen Vibrio Cholerae.” Nature 406 (6795): 477–83.

Jacob, F., and J. Monod. 1961. “On the Regulation of Gene Activity.” Cold Spring Harbor Symposia on Quantitative Biology 26 (0): 193–211.

Jain, Aayushi, Jain Aayushi, and Srivastava Preeti. 2013. “Broad Host Range Plasmids.” FEMS Microbiology Letters 348 (2): 87–96.

Katashkina, Joanna I., Hara Yoshihiko, Lyubov I. Golubeva, Irina G. Andreeva, Tatiana M. Kuvaeva, and Sergey V. Mashko 2009. “Use of the λ Red-Recombineering Method for Genetic Engineering of Pantoea Ananatis.” BMC Molecular Biology 10 (1): 34.

Keseler, Ingrid M., Amanda Mackie, Martin Peralta-Gil, Alberto Santos-Zavaleta, Socorro Gama-Castro, Cesar Bonavides-Martmez, Carol Fulcher, et al. 2013. “EcoCyc: Fusing Model Organism Databases with Systems Biology.” Nucleic Acids Research 41 (Database issue): D605–12.

Lajoie, Marc J., Alexis J. Rovner, Daniel B. Goodman, Hans-Rudolf Aerni, Adrian D. Haimovich, Gleb Kuznetsov, Jaron A. Mercer, et al. 2013. “Genomically Recoded Organisms Expand Biological Functions.” Science 342 (6156): 357–60.

Larson, Matthew H., Luke A. Gilbert, Wang Xiaowo, Wendell A. Lim, Jonathan S. Weissman, and Lei S. Qi. 2013. “CRISPR Interference (CRISPRi) for Sequence-Specific Control of Gene Expression.” Nature Protocols 8 (11): 2180–96.

Li, Mamie Z., and Stephen J. Elledge. 2007. “Harnessing Homologous Recombination in Vitro to Generate Recombinant DNA via SLIC.” Nature Methods 4 (3): 251–56.

Litcofsky, Kevin D., Raffi B. Afeyan, Russell J. Krom, Ahmad S. Khalil, and James J. Collins. 2012. “Iterative Plug-and-Play Methodology for Constructing and Modifying Synthetic Gene Networks.” Nature Methods 9 (11): 1077–80.

Lutz, R., and H. Bujard. 1997. “Independent and Tight Regulation of Transcriptional Units in Escherichia Coli via the LacR/O, the TetR/O and AraC/I1-I2 Regulatory Elements.” Nucleic Acids Research 25 (6): 1203–10.

Maida, I., E. Bosi, E. Perrin, M. C. Papaleo, V. Orlandini, M. Fondi, R. Fani, J. Wiegel, G. Bianconi, and F. Canganella. 2013. “Draft Genome Sequence of the Fast-Growing Bacterium Vibrio Natriegens Strain DSMZ 759.” Genome Announcements 1 (4): e00648-13 – e00648-13.

Messing, J., B. Gronenborn, B. Müller-Hill, and P. Hans Hopschneider. 1977. “Filamentous Coliphage M13 as a Cloning Vehicle: Insertion of a HindII Fragment of the Lac Regulatory Region in M13 Replicative Form in Vitro.” Proceedings of the National Academy of Sciences of the United States of America 74 (9): 3642–46.

Overbeek, Ross, Overbeek Ross, Olson Robert, Gordon D. Pusch, Gary J. Olsen, James J. Davis, Disz Terry, et al. 2013. “The SEED and the Rapid Annotation of Microbial Genomes Using Subsystems Technology (RAST).” Nucleic Acids Research 42 (D1): D206–14.

Payne, W. J., R. G. Eagon, and A. K. Williams. 1961. “Some Observations on the Physiology of Pseudomonas Natriegens Nov. Spec.” Antonie van Leeuwenhoek 27: 121–28.

Peters, Jason M., Alexandre Colavin, Handuo Shi, Tomasz L. Czarny, Matthew H. Larson, Spencer Wong, John S. Hawkins, et al. 2016. “A Comprehensive, CRISPR-Based Functional Analysis of Essential Genes in Bacteria.” Cell 165 (6): 1493–1506.

Qi, Lei S., Matthew H. Larson, Luke A. Gilbert, Jennifer A. Doudna, Jonathan S. Weissman, Adam P. Arkin, and Wendell A. Lim. 2013. “Repurposing CRISPR as an RNA-Guided Platform for Sequence-Specific Control of Gene Expression.” Cell 152 (5): 1173–83.

Sabri, Suriana, Lars K. Nielsen, and Claudia E. Vickers. 2013. “Molecular Control of Sucrose Utilization in Escherichia Coli W, an Efficient Sucrose-Utilizing Strain.” Applied and Environmental Microbiology 79 (2): 478–87.

Schleif, R. 2000. “Regulation of the L-Arabinose Operon of Escherichia Coli.” Trends in Genetics: TIG 16 (12): 559–65.

Shimizu, Tohru, Kaori Ohtani, Hideki Hirakawa, Kenshiro Ohshima, Atsushi Yamashita, Tadayoshi Shiba, Naotake Ogasawara, Masahira Hattori, Satoru Kuhara, and Hideo Hayashi. 2002. “Complete Genome Sequence of Clostridium Perfringens, an Anaerobic Flesh-Eater.” Proceedings of the National Academy of Sciences of the United States of America 99 (2): 996–1001.

van Opijnen, Tim, and Andrew Camilli. 2010. “Genome-Wide Fitness and Genetic Interactions Determined by Tn-Seq, a High-Throughput Massively Parallel Sequencing Method for Microorganisms.” Vurrent Protocols in Microbiology Chapter 1 (November): Unit1E.3.

Waldor, M. K., and J. J. Mekalanos. 1996. “Lysogenic Conversion by a Filamentous Phage Encoding Cholera Toxin.” Science 272 (5270): 1910–14.

Wang, Harris H., Farren J. Isaacs, Peter A. Carr, Zachary Z. Sun, George Xu, Craig R. Forest, and George M. Church. 2009. “Programming Cells by Multiplex Genome Engineering and Accelerated Evolution.” Nature 460 (7257): 894–98.

Wang, Z., B. Lin, W. J. Hervey, and G. J. Vora. 2013. “Draft Genome Sequence of the Fast-Growing Marine Bacterium Vibrio Natriegens Strain ATCC 14048.” Genome Announcements 1 (4): e00589-13 – e00589-13.

Willardsen, R. R., Willardsen R.R., F. F. Busta, C. E. Allen, and L. B. Smith. 1978. “GROWTH AND SURVIVAL OF Clostridium Perfringens DURING CONSTANTLY RISING TEMPERATURES.” Journal of Food Science 43 (2): 470–75.

## References

Aziz, Ramy K., Daniela Bartels, Aaron A. Best, Matthew DeJongh, Terrence Disz, Robert A. Edwards, Kevin Formsma, et al. 2008. “The RAST Server: Rapid Annotations Using Subsystems Technology.” BMC Genomics 9 (February): 75.

Cookson, Scott, Natalie Ostroff, Wyming Lee Pang, Dmitri Volfson, and Jeff Hasty. 2005. “Monitoring Dynamics of Single-Cell Gene Expression over Multiple Cell Cycles.” Molecular Systems Biology 1 (November): 2005.0024.

Duffy, D. C., J. C. McDonald, O. J. Schueller, and G. M. Whitesides. 1998. “Rapid Prototyping of Microfluidic Systems in Poly(dimethylsiloxane).” Analytical Chemistry 70 (23): 4974–84.

Gibson, Daniel, and Gibson Daniel. 2009. “One-Step Enzymatic Assembly of DNA Molecules up to Several Hundred Kilobases in Size.” Protocol Exchange. doi:10.1038/nprot.2009.77.

Katashkina, Joanna I., Hara Yoshihiko, Lyubov I. Golubeva, Irina G. Andreeva, Tatiana M. Kuvaeva, and Sergey V. Mashko. 2009. “Use of the λ Red-Recombineering Method for Genetic Engineering of Pantoea Ananatis.” BMC Molecular Biology 10 (1): 34.

Krumsiek, Jan, Roland Arnold, and Thomas Rattei. 2007. “Gepard: A Rapid and Sensitive Tool for Creating Dotplots on Genome Scale.” Bioinformatics 23 (8): 1026–28.

Langmead, Ben, Cole Trapnell, Mihai Pop, and Steven L. Salzberg. 2009. “Ultrafast and Memory-Efficient Alignment of Short DNA Sequences to the Human Genome.” Genome Biology 10 (3): R25.

Martínez-García, Esteban, Belén Calles, Miguel Arévalo-Rodríguez, and Víctor de Lorenzo. 2011. “pBAM1: An All-Synthetic Genetic Tool for Analysis and Construction of Complex Bacterial Phenotypes.” BMC Microbiology 11 (February): 38.

Stricker, Jesse, Scott Cookson, Matthew R. Bennett, William H. Mather, Lev S. Tsimring, and Jeff Hasty. 2008. “A Fast, Robust and Tunable Synthetic Gene Oscillator.” Nature 456 (7221): 516–19.

Takahashi, Chris N., Aaron W. Miller, Felix Ekness, Maitreya J. Dunham, and Eric Klavins. 2015. “A Low Cost, Customizable Turbidostat for Use in Synthetic Circuit Characterization.” ACS Synthetic Biology 4 (1): 32–38.

van Opijnen, Tim, and Andrew Camilli. 2010. “Genome-Wide Fitness and Genetic Interactions Determined by Tn-Seq, a High-Throughput Massively Parallel Sequencing Method for Microorganisms.” Current Protocols in Microbiology Chapter 1 (November): Unit1E.3.

Vega, Nicole M., Kyle R. Allison, Ahmad S. Khalil, and James J. Collins. 2012. “Signaling-Mediated Bacterial Persister Formation.” Nature Chemical Biology 8 (5): 431–33.

